# Distinct viability and oxygen-sensing responses to hypoxia and reoxygenation in pancreatic islets

**DOI:** 10.64898/2026.10.18.759980

**Authors:** Hiroyuki Kato, Heather N. Zook, Neslihan Erdem, Hiroaki Mitsugashira, Hsun Teresa Ku, Hirotake Komatsu

## Abstract

Early graft hypoxia limits the efficiency and success of islet transplantation. Defining the relationships among oxygen availability, exposure duration, islet survival, and oxygen sensing is therefore critical, but these relationships remain incompletely understood. We examined adult primary rat islets under continuous exposure to 1%, 5%, or 18% O_2_ for 48 or 96 h and under reoxygenation conditions comprising 48 h at 1% O_2_ followed by 48 h at 18% O_2_. We assessed islet size-dependent viability, morphology, mass recovery, estimated viable mass, and targeted hypoxia-responsive gene-expression signatures. Overall, 6,699 individual islets were analyzed. At 1% O_2_, viability decreased significantly, with most loss occurring within the first 48 h (93.0% at 0 h, 70.8% at 48 h, and 65.0% at 96 h). Overall viability remained largely preserved at 5% and 18% O_2_ (92.8 and 95.0% at 96h, respectively), although larger islets showed vulnerability at 5% O_2_. Exposure to 1% O_2_ caused deformation, particularly in larger islets, and reduced estimated viable mass compared with 5% and 18% O_2_. Reoxygenation did not fully restore viability or estimated viable mass (76.0% at 96h). Gene-expression analyses revealed sustained hypoxia-responsive signatures at both 1% and 5% O_2_, distinct from those at 18% O_2_. Reoxygenation shifted gene-expression profiles toward a normoxia-like signature. These findings suggest divergent viability and oxygen-sensing responses to hypoxia and reoxygenation. The 5% O_2_ condition may represent a marginal oxygen environment that preserves short-term overall survival despite sustained molecular responses to hypoxia. These data inform the oxygen targets and timing of early graft oxygenation strategies.

## INTRODUCTION

Pancreatic islets are highly vulnerable to oxygen limitation due to their cellular oxygen demand and structure (1–3). Their three-dimensional architecture creates oxygen gradients from the surface toward the core, while relatively high metabolic demand further contributes to oxygen depletion (4). Compared to smaller islets, larger islets are particularly susceptible because oxygen must diffuse over greater distances to reach centrally located cells, increasing the risk of central necrosis. This problem becomes especially important immediately after transplantation, before new vascular connections are established (5). Substantial islet loss occurs during this early period, partly because of the harsh graft environment, including severe hypoxia. Consequently, early graft survival is influenced by the severity and duration of hypoxia, as well as islet size and cellular oxygen demand (4). However, islet responses to varying oxygen conditions remain incompletely characterized.

Clinically, islets have been transplanted into the portal vein, where they engraft throughout the liver parenchyma in a dispersed pattern. More recently, extrahepatic transplantation sites have gained interest, particularly for stem cell-derived grafts, because they offer improved access for graft monitoring and retrieval (6). These sites may also benefit conventional transplantation of donor-derived islets because they allow less invasive intervention than intrahepatic transplantation; clinical studies have explored both subcutaneous and intramuscular sites (7). However, limited oxygen availability remains a major challenge due to the high graft packing density at these sites. Before effective revascularization develops, transplanted cells depend primarily on oxygen diffusion from surrounding tissues (3, 4, 8). Densely packed grafts may therefore develop substantial intragraft hypoxia. Furthermore, graft oxygen conditions change over time after transplantation according to surrounding tissue oxygen levels, graft density, and revascularization status.

The goal of the present study was to characterize islet survival phenotypes under various oxygen conditions, as well as oxygen-sensing responses at the gene-expression level. These findings will help guide transplantation strategies, including oxygenation interventions. We examined these responses over exposure durations relevant to the early post-transplant period. We used an oxygen-permeable culture platform to expose adult primary rat islets to precisely controlled 1% O_2_ (severe hypoxia), 5% O_2_ (a physiologically relevant tissue oxygen level), and 18% O_2_ (a well-oxygenated culture condition) for 48 or 96 h (9). Additionally, we model reoxygenation by restoring oxygen levels to 18% O_2_ after 48 h at 1% O_2_. We assessed viability, size-dependent vulnerability, morphology, mass recovery, estimated viable mass, and hypoxia-responsive gene expression to characterize oxygen- and time-dependent responses relevant to early graft preservation.

## Materials and Methods

### Isolation and culture of primary adult rat islets

Adult islets were isolated from 16–20-week-old wild-type female Lewis rats using our previously described procedure (9–11). Briefly, pancreata were distended with collagenase through the common bile duct, enzymatically digested, purified by Histopaque-1077 density-gradient centrifugation, and hand-picked. Islet yield was quantified as islet equivalents (IEQ), with a 150-µm-diameter islet defined as 1 IEQ (12). Islets were recovered overnight at 27°C in supplemented CMRL 1066 medium, before the subsequent experiments.

### Islet culture under controlled oxygen conditions

Oxygen-permeable culture plates with a micropyramid surface architecture were used to provide well-defined oxygen environments surrounding individual spheroids, as previously described (9, 13). Cultures were maintained at 1% or 5% O_2_ using controlled-O_2_ chambers placed in a 37°C incubator, or at approximately 18% O_2_ in a conventional humidified incubator supplied with 5% CO_2_ and air, accounting for the partial pressure of water vapor (13). Approximately 200 IEQ were cultured per well in a 24-well format for 48 or 96 h under continuous O_2_ conditions. For the reoxygenation experiments, samples were cultured at 1% O_2_ for the first 48 h and then transferred to 18% O_2_ for an additional 48 h. The experimental protocol is summarized in **Fig. 1**.

**Figure 1.**
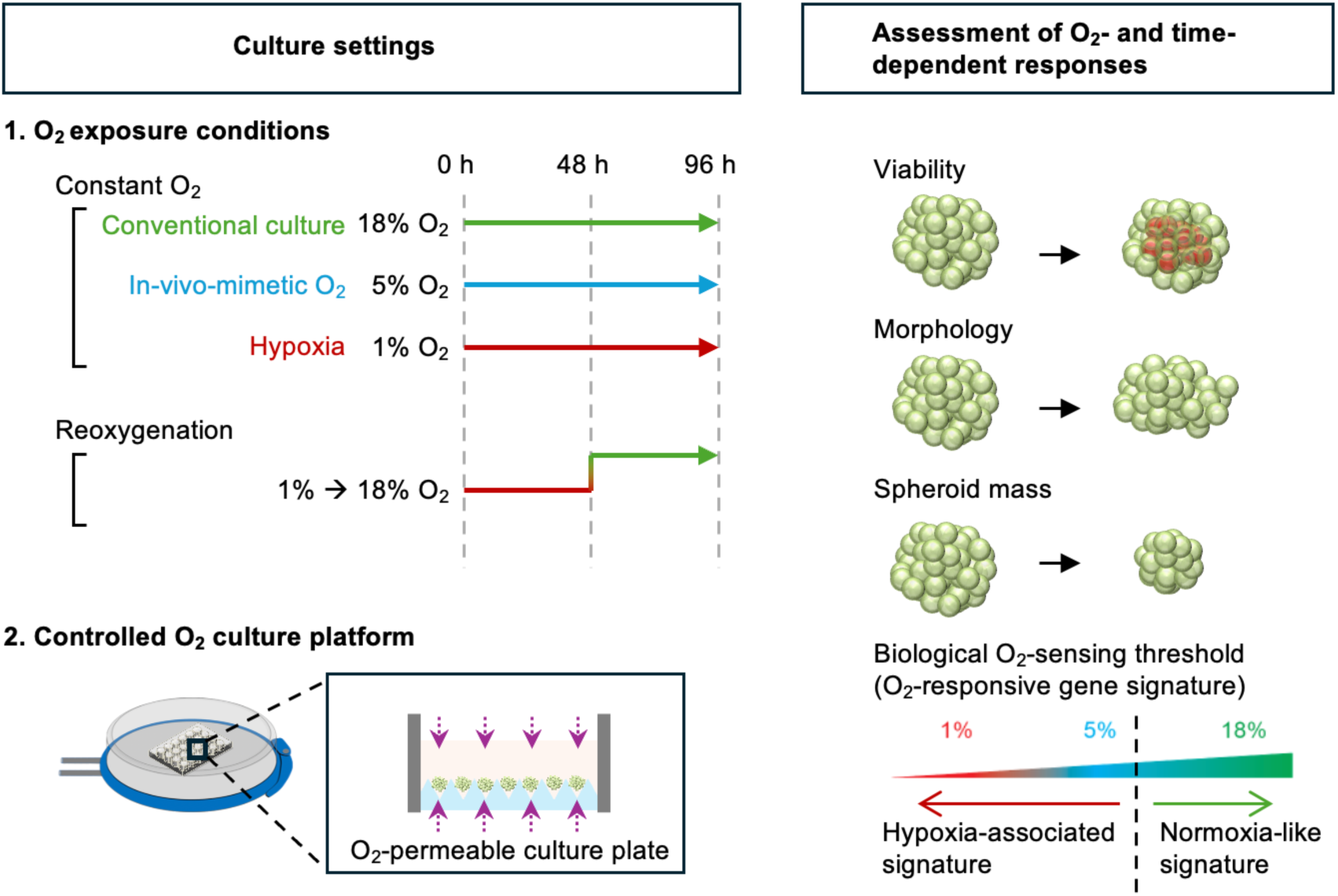
Experimental design for assessment of O_2_- and time-dependent responses in adult primary rat islets. Adult primary rat islets were cultured using an O_2_-permeable culture platform under continuous exposure to 1%, 5%, or 18% O_2_ for 48 or 96 h. For reoxygenation studies, islets were cultured at 1% O_2_ for the first 48 h and then transferred to 18% O_2_ for an additional 48 h. The 18% O_2_ condition represents conventional culture, 5% O_2_ represents a physiologically relevant tissue oxygen level, and 1% O_2_ represents severe hypoxia. O_2_- and time-dependent responses were assessed by measuring viability, morphology, islet mass recovery, estimated viable mass, and O_2_-responsive gene-expression signatures. The lower-right schematic conceptually illustrates oxygen sensing assessed through gene-expression responses, ranging from a hypoxia-associated to a normoxia-like signature; it does not indicate a precisely determined oxygen-sensing threshold.

### Quantitative viability assessment, spheroid/islet size and morphology

We assessed individual islet viability by live/dead staining using our previously reported method (9, 13, 14). Briefly, spheroids were stained with 0.48 µM FDA and 15 µM PI for 5 min at room temperature, imaged by wide-field fluorescence microscopy (Leica, Wetzlar, Germany), and analyzed using cellSens software (EVIDENT, Tokyo, Japan) to calculate area-based viability from live and dead imaging. Islet size (diameter) was estimated from two-dimensional projected area as previously described (15).

Morphology was quantified using shape factor, which is calculated from the area and perimeter of individual spheroids/islets as previously described (9). Shape factor ranges from 0 to 1, with values approaching 1 indicating a more circular morphology.

### Gene-expression analysis by microfluidic quantitative PCR

Gene-expression analysis was performed as previously described (16). Total RNA was isolated from cultured primary rat islets, and cDNA was synthesized. Gene expression was quantified using the BioMark 48.48 Dynamic Array platform (Standard BioTools, South San Francisco, CA) with gene-specific TaqMan assays listed in **Supplementary Table 1**. Target-gene expression was normalized to *beta actin (Actb)*. A 27-gene panel comprising known hypoxia-related genes across several biological categories, including oxygen sensing, stress signaling, metabolic adaptation, vascular/matrix remodeling, and immune/inflammatory responses, was used for subsequent analyses. *Actb*-normalized expression values were log2-transformed for heatmap visualization and principal component analysis (PCA).

### Statistical analysis

Data are presented as mean ± SEM, with individual biological experiments or preparations shown where applicable. Differences among experimental conditions were evaluated by one-way ANOVA followed by Tukey-Kramer multiple-comparison testing. Size-stratified viability and morphology analyses were performed within predefined size categories. Linear regression was used descriptively to assess associations between spheroid/islet diameter and viability, with slopes and R2 values reported. Gene-expression measurements and PCA-derived PC1 and PC2 scores were analyzed by one-way ANOVA with Tukey-Kramer testing, as applicable. Analyses were performed using JMP 19 (SAS Institute Inc.). P < 0.05 was considered statistically significant.

## RESULTS

### Severe hypoxia markedly reduces viability of adult primary rat islets

We analyzed 6,699 adult primary rat islets from four independent isolation preparations across pre-culture, continuous oxygen-exposure, and reoxygenation conditions. Sample numbers are summarized in **Supplementary Table 2**, and overall and preparation-specific size distributions are shown in **Supplementary Fig. S1**. Representative live/dead images showed prominent dead-cell staining under 1% O_2_ at both 48 and 96 h, particularly within central islet regions, whereas islets cultured at 5% and 18% O_2_ remained predominantly viable **(Fig. 2A)**.

**Figure 2.**
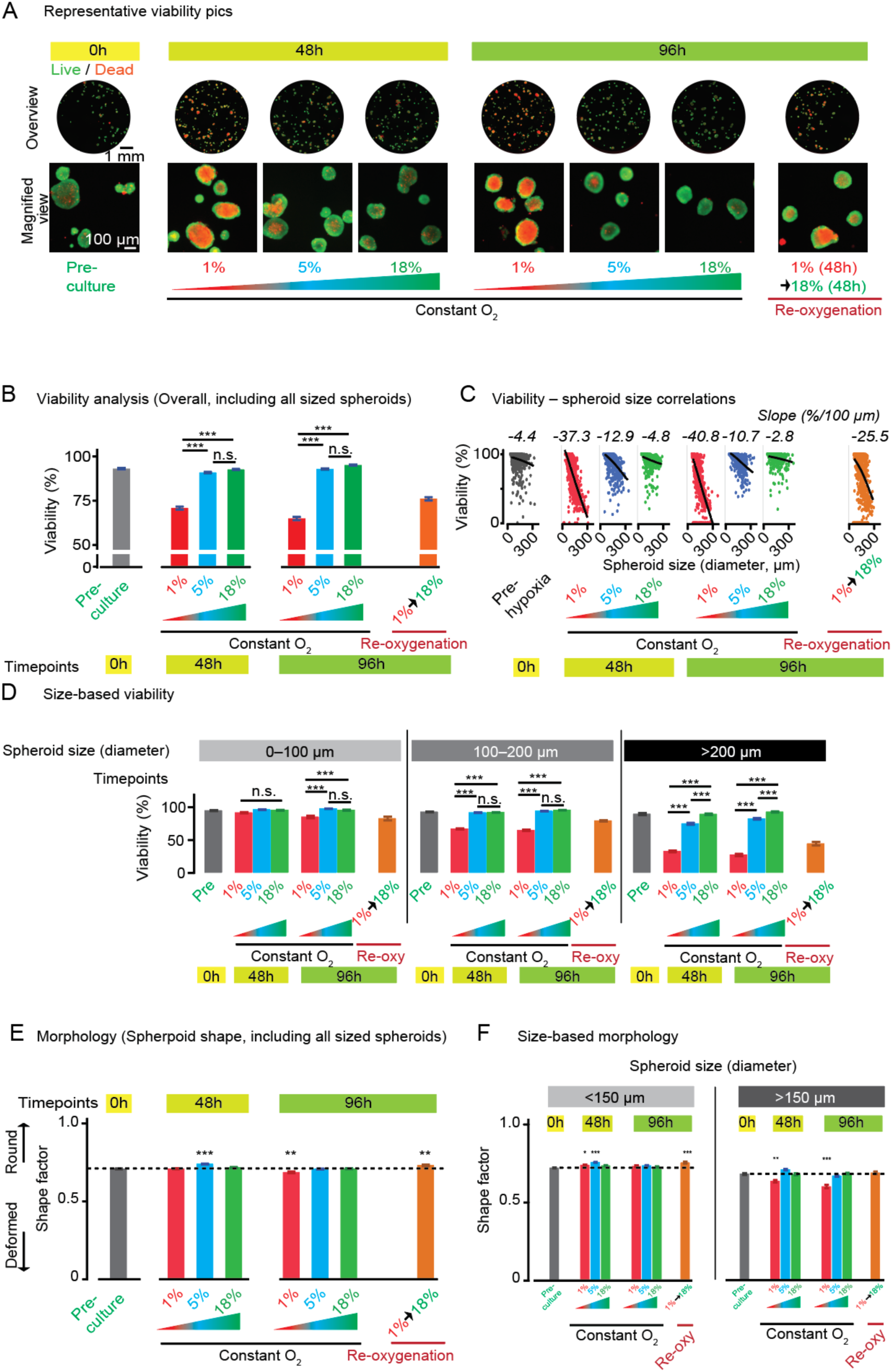
Viability of adult primary rat islets under different oxygen conditions. **(A)** Representative live/dead fluorescence images of adult primary rat islets before hypoxic exposure (0 h) and following 48 or 96 h of culture under the indicated oxygen conditions. At 48 h, islets were cultured continuously at 1%, 5%, or 18% O_2_. At 96 h, samples included continuous exposure to 1%, 5%, or 18% O_2_ and reoxygenation conditions consisting of 48 h at 1% O_2_ followed by 48 h at 18% O_2_. Overview and higher-magnification images are shown. Green and red fluorescence indicate live and dead cells, respectively. Scale bars, 1 mm for overview images and 100 µm for magnified images. **(B)** Overall islet viability quantified across all islet sizes under the indicated culture conditions. A total of 6,699 islets from four independent adult primary rat-islet preparations were included in the analysis. Bars represent mean ± SEM. Differences among experimental conditions were evaluated by one-way ANOVA followed by Tukey– Kramer multiple-comparison testing. Selected comparisons among 1%, 5%, and 18% O_2_ conditions at 48 and 96 h are indicated in the panel; ***P < 0.0001; n.s., not significant. The number of islets analyzed under each condition is provided in **(Supplementary Table 2)**. Overall and preparation-specific islet-size distributions are shown in **(Supplementary Fig. S1)**, and full spheroid-level viability distributions, including preparation-specific distributions, are shown in **(Supplementary Fig. S2)**. Complete Tukey–Kramer pairwise comparisons, including reoxygenation conditions, are provided in **(Supplementary Table 3)**. **(C)** Relationship between islet diameter and viability in adult primary rat islets under the indicated oxygen conditions. Individual islet-level measurements from four independent preparations were pooled within each condition, and simple linear regression was used to describe the association between islet size and viability. Regression slopes are shown above each condition and are expressed as the change in viability percentage points per 100 µm increase in islet diameter. Pre-hypoxia, continuous 1%, 5%, and 18% O_2_ conditions at 48 and 96 h, and reoxygenation condition. The regression analyses were used descriptively rather than for formal between-condition statistical comparisons. Full regression parameters, including n, slope, and R2 values, are provided in **(Supplementary Table 4)**. (D) Size-stratified viability analysis of adult primary rat islets. Islets were categorized by diameter as 0–100 µm, 100–200 µm, or >200 µm, and viability was analyzed separately within each size category under the indicated oxygen conditions. Data include pre-hypoxia, continuous culture at 1%, 5%, or 18% O_2_ for 48 or 96 h, and reoxygenation condition. Measurements from four independent islet preparations were pooled within each condition and size category. Bars represent mean ± SEM. Differences among oxygen conditions within each size category were evaluated by one-way ANOVA followed by Tukey–Kramer multiple-comparison testing. Selected pairwise comparisons are indicated in the panel; ***P < 0.0001 and n.s., not significant. Complete sample numbers and pairwise comparisons for each size category are provided in (**Supplementary Table 5**). **(E) Morphological analysis of adult primary rat islets under different oxygen conditions.** Overall islet morphology quantified by shape factor across all islet sizes before hypoxic exposure and following 48 or 96 h under continuous oxygen-exposure or the reoxygenation condition. Shape-factor values approaching 1 indicate a more rounded morphology, whereas lower values indicate increasing deformation. Bars represent mean ± SEM. Differences among experimental conditions were evaluated by one-way ANOVA followed by Tukey–Kramer multiple-comparison testing. Asterisks indicate selected comparisons with the pre-hypoxia condition (**\*\*\*P < 0.0001; **P < 0.01**). Complete Tukey–Kramer pairwise comparisons are provided in **(Supplementary Table 6)**. **(F)** Size-stratified morphology analysis of adult primary rat islets. Islets were categorized by diameter as ≤150 µm or >150 µm, and shape factor was analyzed separately within each size category under the indicated oxygen conditions. Data include pre-hypoxia, continuous culture at 1%, 5%, or 18% O_2_ for 48 or 96 h, and the reoxygenation condition. Measurements from four independent islet preparations were pooled within each condition and size category. Bars represent mean ± SEM. Differences among experimental conditions within each size category were evaluated by one-way ANOVA followed by Tukey–Kramer multiple-comparison testing. Selected comparisons are indicated in the panel; ***P < 0.0001, **P < 0.01, and *P < 0.05. Complete sample numbers and pairwise comparisons for each size category are provided in **(Supplementary Table 7)**.

Quantitative analysis confirmed a pronounced oxygen-dependent effect **(Fig. 2B)**. Mean viability decreased from 93.0% pre-culture to 70.8% and 65.0% at 1% O_2_ after 48 and 96 h, respectively. Thus, viability loss occurred predominantly within the first 48 h, followed by a modest further decline by 96 h. At both timepoints, viability at 1% O_2_ was significantly lower than at 5% and 18% O_2_ (all P < 0.0001), whereas viability at 5% and 18% O_2_ remained statistically similar. Reoxygenation did not fully restore viability to that observed under the corresponding continuous higher-O_2_ condition. Full viability distributions of individual islets are shown in **Supplementary Fig. S2**, and complete Tukey-Kramer comparisons are provided in **Supplementary Table 3**.

### Islet size is strongly associated with hypoxia-dependent viability loss

Pooled islet-level regression revealed a marked size-dependent decline in viability under severe hypoxia **(Fig. 2C)**. At 48 h, the slope was -37.3 percentage points per 100 µm at 1% O_2_, compared with -12.9 at 5% and -4.8 at 18% O_2_; a similar pattern persisted at 96 h. Reoxygenated islets also showed substantial size-dependent viability loss.

Thus, larger adult islets were particularly vulnerable to severe hypoxia, whereas size had relatively little association with viability under continuous 18% O_2_. Full regression parameters are provided in **Supplementary Table 4**.

### Size stratification reveals pronounced oxygen sensitivity in larger adult islets

To further examine this relationship, islets were stratified into 0–100 µm, 100–200 µm, and >200 µm groups **(Fig. 2D)**. The smallest islets showed relatively limited oxygen-dependent differences, whereas viability loss at 1% O_2_ became increasingly pronounced with larger islet size. In the >200 µm group, viability was markedly reduced at 1% O_2_ and was also significantly lower at 5% than at 18% O_2_ at both 48 and 96 h.

This graded response was not apparent in the overall viability analysis and was consistent with the regression findings. Complete size-stratified comparisons are provided in **Supplementary Table 5**.

### Adult primary rat islets largely preserve morphology across oxygen conditions

We quantified morphology using shape factor **(Fig. 2E)**. Adult rat islets largely maintained their overall roundness across oxygen conditions, despite significant differences in viability. Prolonged exposure to 1% O_2_ produced only a modest reduction in shape factor, whereas 5% and 18% O_2_ remained close to pre-culture morphology.

Reoxygenation also preserved overall islet shape. **Supplementary Table 6** provides the complete Tukey-Kramer comparisons.

### Morphological deformation is concentrated in larger adult islets

We stratified adult islets by size to determine whether diameter influenced morphological responses **(Fig. 2F)**. Islets ≤150 µm generally maintained or modestly increased their roundness across culture conditions. In contrast, islets >150 µm developed selective deformation under 1% O_2_, with the reduction in shape factor becoming more pronounced by 96 h, while 5% and 18% O_2_ remained near pre-culture levels. Reoxygenated larger islets also maintained morphology similar to pre-culture. Overall, sustained severe hypoxia predominantly affected the morphology of larger adult islets. **Supplementary Table 7** provides complete sample numbers and pairwise comparisons.

### Adult primary rat islets largely maintain mass during culture

We compared corresponding pre- and post-culture images to assess changes in adult primary rat-islet mass **(Fig. 3A)**. Representative images showed relatively limited changes in islet mass, estimated by area-based image analysis, across oxygen conditions and culture durations. For imaging, we collected all islets dispersed within each opaque micropyramid-bottomed culture well, transferred them to a flat, translucent plate, and gathered them at the center with minimal spacing to capture the entire preparation.

**Figure 3.**
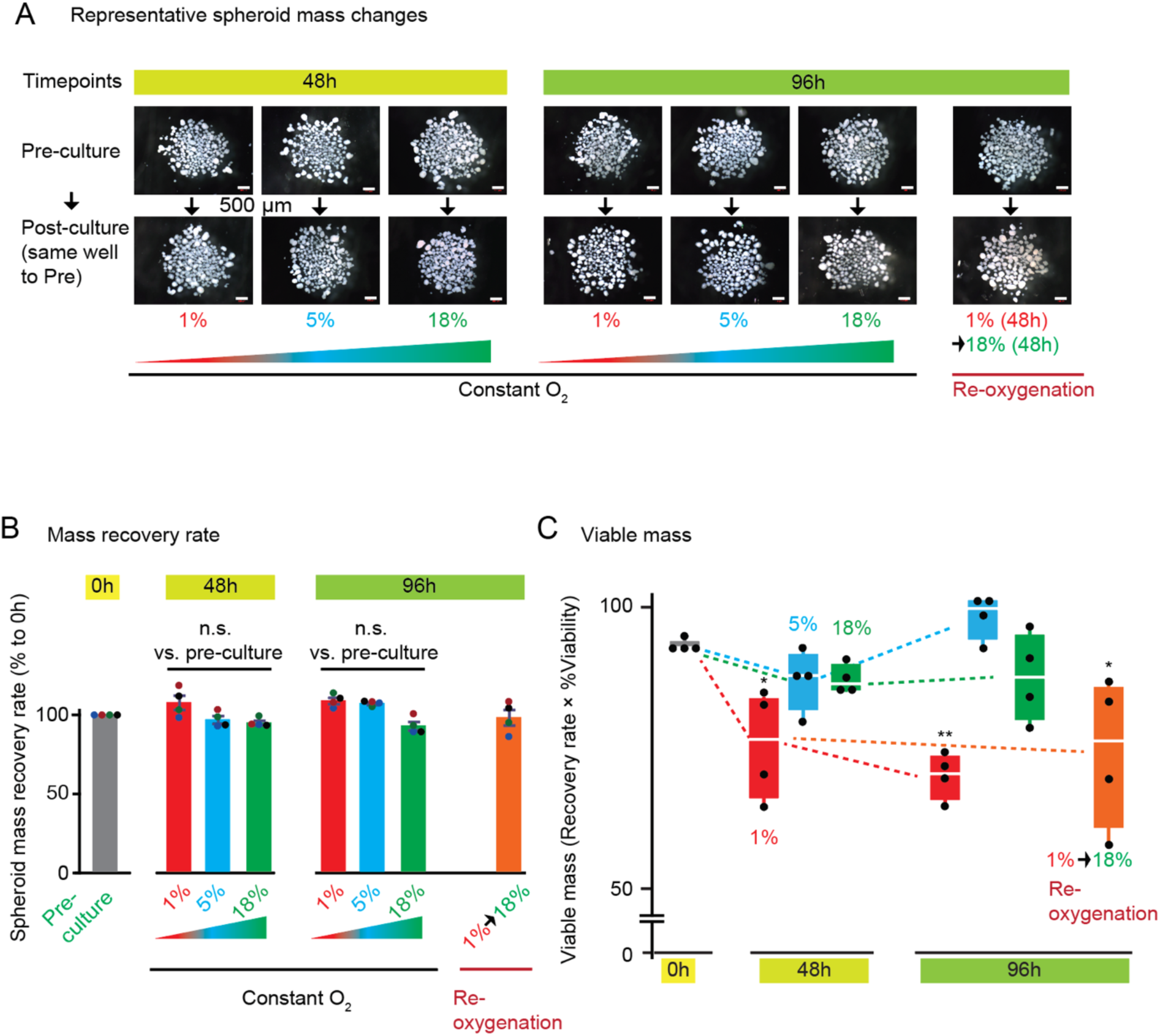
Assessment of mass recovery in adult primary rat islets under different oxygen conditions. **(A)** Representative images used to assess changes in adult primary rat-islet mass before and after culture. Islets were transferred to multiwell plates for standardized imaging, and corresponding wells were imaged before culture and after 48 or 96 h under the indicated oxygen conditions. The upper row shows pre-culture images and the lower row shows the corresponding post-culture preparations. At 48 h, islets were cultured continuously at 1%, 5%, or 18% O_2_. At 96 h, conditions included continuous exposure to 1%, 5%, or 18% O_2_ and reoxygenation consisting of 48 h at 1% O_2_ followed by 48 h at 18% O_2_. Images are representative of four independent islet preparations. Scale bar, 500 µm. **(B)** Quantification of islet mass recovery. Islet mass was estimated from two-dimensional islet area measured from corresponding pre- and post-culture images. Mass recovery rate was calculated as post-culture islet area divided by the corresponding pre-culture area and expressed as a percentage of the pre-culture value. Bars represent mean ± SEM from four independent islet preparations; individual points represent independent preparations. Differences among experimental conditions were evaluated by one-way ANOVA followed by Tukey–Kramer multiple-comparison testing. Selected comparisons with the pre-hypoxia condition are indicated in the panel; n.s., not significant. Exact adjusted P values and complete pairwise comparisons are provided in **(Supplementary Table 8)**. **(C)** Quantification of viable mass in adult primary rat islets. Viable mass was calculated for each independent preparation by multiplying the islet mass recovery rate by the corresponding viability value and was expressed relative to the pre-culture measurement. Data include pre-culture, continuous culture at 1%, 5%, or 18% O_2_ for 48 or 96 h, and the reoxygenation condition. Data from four independent islet preparations are shown, with individual points representing independent preparations. Differences among experimental conditions were evaluated by one-way ANOVA followed by Tukey– Kramer multiple-comparison testing. Selected comparisons with the pre-culture condition are indicated in the panel; ***P < 0.0001, **P < 0.01, and *P < 0.05. Exact adjusted P values and complete pairwise comparisons are provided in **(Supplementary Table 9)**.

Quantitative analysis revealed relatively limited changes in islet mass, estimated by area-based image analysis, across oxygen conditions and culture durations **(Fig. 3B)**. Mean mass recovery remained near the pre-culture reference value after both 48 and 96 h, with no significant reduction under 1%, 5%, or 18% O_2_ conditions. Reoxygenation also preserved overall mass. Thus, adult primary rat islets maintain mass during prolonged culture across a broad range of oxygen conditions, despite significant loss of viability under severe hypoxia. Complete Tukey-Kramer comparisons are provided in **Supplementary Table 8**.

### Severe hypoxia reduces viable mass in adult primary rat islets

We calculated estimated viable mass by integrating mass recovery with the corresponding viability measurement **(Fig. 3C)**. After 48 h, viable mass decreased to 75.3% at 1% O_2_, compared with 86.8% at 5% O_2_ and 87.0% at 18% O_2_; the reduction at 1% O_2_ was significant relative to pre-culture (P = 0.0483). At 96 h, estimated viable mass remained lowest at 1% O_2_ (69.8%), whereas values at 5% and 18% O_2_ remained comparable to the pre-culture value. Reoxygenation did not restore estimated viable mass to the levels observed under continuous culture at 5% or 18% O_2_. Overall, sustained 1% O_2_ produced the most consistent reduction in viable mass, while continuous culture at 5% and 18% O_2_ largely preserved viable islet mass. Complete Tukey-Kramer comparisons are provided in **Supplementary Table 9**.

### Adult primary rat islets show a robust and graded oxygen-dependent gene-expression response

We examined a 27-gene hypoxia-related panel in adult primary rat islets across the tested oxygen conditions **(Fig. 4A)**. Overall, the heatmap showed clear gene-expression responses at both 1% and 5% O_2_ relative to 18% O_2_ at both timepoints. These changes were broad and coordinated, although not all hypoxia-related genes were upregulated under reduced oxygen conditions. For example, *Hif1a* did not change significantly. Oxygen availability regulates HIF-1α primarily through protein stability rather than *Hif1a* transcription (17, 18). In contrast, many metabolic adaptation genes downstream of the HIF-1α pathway, such as *Gapdh* and *Ldha*, responded significantly to hypoxia at both 48 and 96 h. These responses remained evident at 96 h without an apparent overall attenuation relative to 48 h, suggesting sustained hypoxia-responsive signaling during prolonged exposure. Reoxygenation following initial exposure to 1% O_2_ shifted the gene-expression profile toward that observed under continuous culture at 18% O_2_.

**Figure 4.**
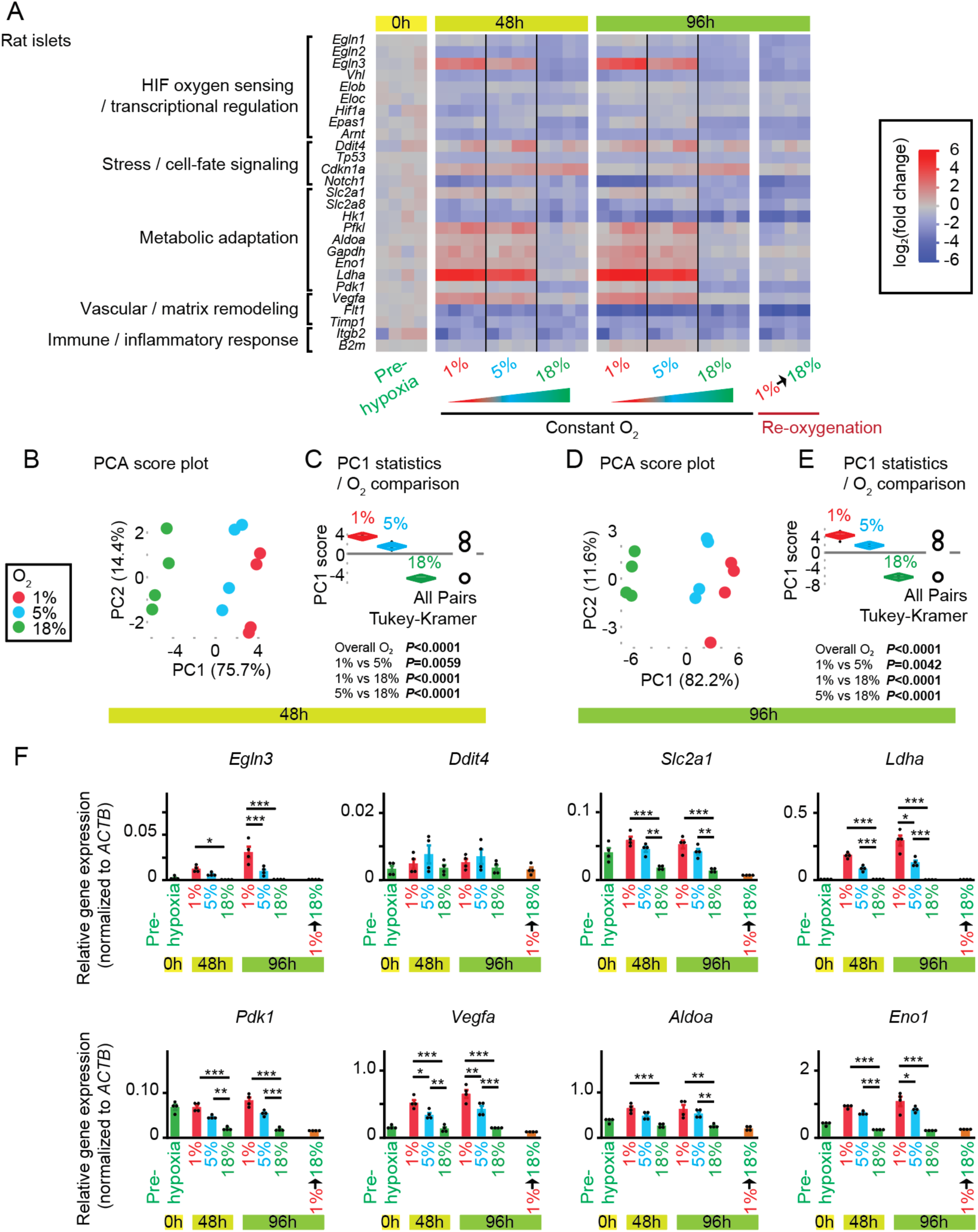
Oxygen-dependent transcriptional responses in adult primary rat islets. **(A)** Heatmap of a 27-gene oxygen-response panel selected primarily based on the KEGG HIF-1 signaling pathway map and grouped into HIF oxygen sensing/transcriptional regulation, stress/cell-fate signaling, metabolic adaptation, vascular/matrix remodeling, and immune/inflammatory response categories. Gene-expression profiles are shown before hypoxic exposure (0 h), after 48 or 96 h of continuous culture at 1%, 5%, or 18% O_2_, and after reoxygenation following 48 h at 1% O_2_ followed by 48 h at 18% O_2_. Colors indicate log2(fold change) according to the scale shown. **(B)** PCA score plot of the 27-gene panel after 48 h of continuous culture at 1%, 5%, or 18% O_2_. Four independent samples were included per oxygen condition. Percentages shown on the axes indicate the variance explained by PC1 and PC2. Detailed statistical results for the PCA are provided in **Supplementary Table 10**. **(C)** Distribution of PC1 scores at 48 h. Overall differences among oxygen conditions were evaluated by one-way ANOVA followed by Tukey–Kramer multiple-comparison testing; corresponding P values are shown in the panel. **(D)** PCA score plot of the same 27-gene panel after 96 h of continuous culture at 1%, 5%, or 18% O_2_. **(E)** Distribution of PC1 scores at 96 h, analyzed by one-way ANOVA followed by Tukey–Kramer multiple-comparison testing. **(F)** Relative expression of eight representative genes selected from the functional categories shown in panel A: *Egln3* (HIF oxygen sensing/transcriptional regulation), *Ddit4* (stress/cell-fate signaling), *Slc2a1*, *Ldha*, *Pdk1*, *Aldoa*, and *Eno1* (metabolic adaptation), and *Vegfa* (vascular/matrix remodeling). Expression values were normalized to *Actb*. Individual samples are shown as points, and bars represent mean ± SEM. Detailed statistical results are provided in **Supplementary Table 11**.

Overall, adult primary rat islets exhibited broadly similar hypoxia-associated gene-expression patterns at 1% and 5% O_2_ that were distinct from those at 18% O_2_.

### PCA further distinguishes 1%, 5%, and 18% O_2_ gene-expression states in adult rat islets

We next used PCA to evaluate overall gene-expression patterns across the 27-gene panel under the three continuous O_2_ conditions **(Fig. 4B–E)**. At 48 h, PC1 captured 75.7% of the total variance and clearly ordered the three oxygen conditions, with 1% and 18% O_2_ at opposite ends and 5% O_2_ occupying an intermediate position closer to 1% O_2_ **(Fig. 4B, C)**. Although 18% O_2_ appeared most clearly separated from the 1% and 5% O_2_ groups, statistical analysis showed significant differences in PC1 scores among all three oxygen conditions, including between 1% and 5% O_2_, with 5% O_2_ occupying an intermediate position. The same pattern persisted at 96 h, when PC1 captured 82.2% of the variance and again significantly separated all three oxygen conditions **(Fig. 4D, E)**. PC2 showed no significant oxygen-dependent separation at either timepoint. PC1 loading analyses showed that genes across oxygen-sensing, stress, metabolic, and vascular pathways contributed to these patterns **(Supplementary Fig. S3A, B)**. Overall, adult rat islets showed a robust and graded oxygen-dependent gene-expression response, with 5% O_2_ consistently occupying an intermediate position along PC1 but showing a profile closer to 1% O_2_ than to 18% O_2_ at both 48 and 96 h. Detailed PCA statistics, including variance explained and PC1 score comparisons among oxygen conditions, are provided in **Supplementary Table 10**.

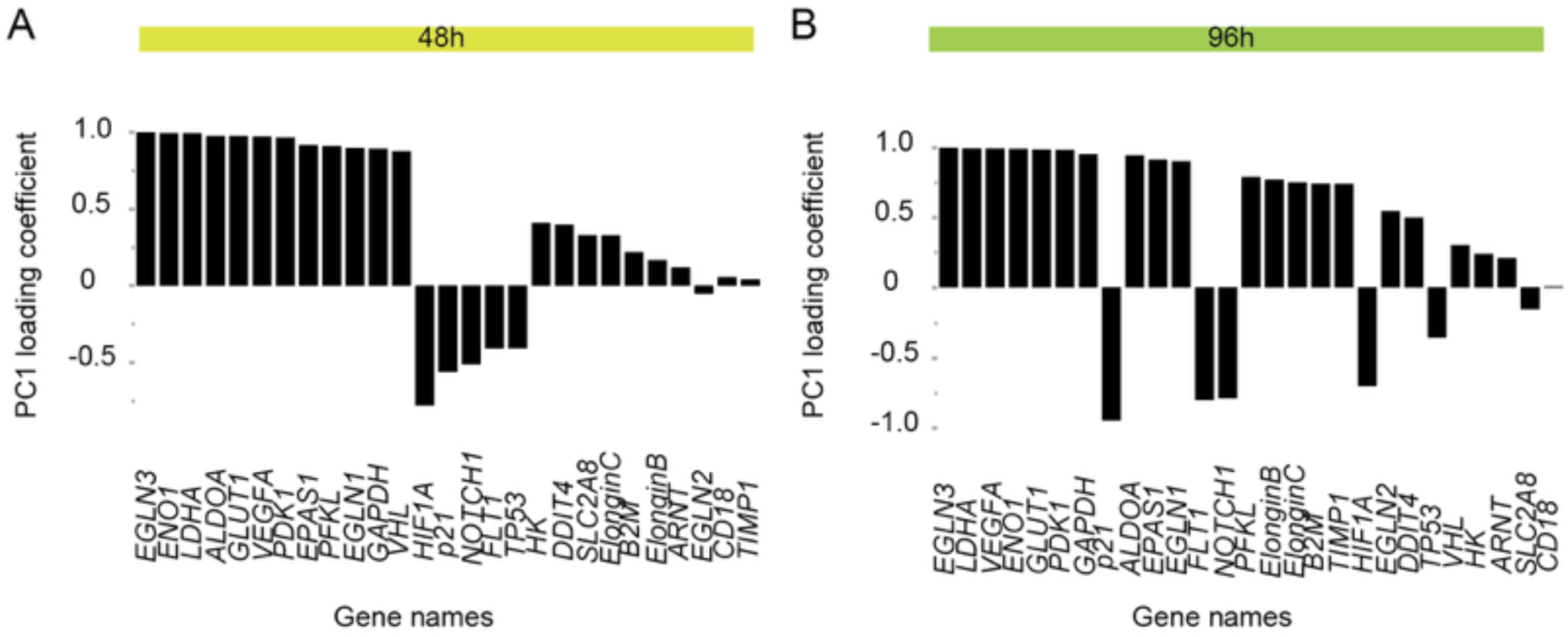

### Representative genes demonstrate a broader oxygen-dependent gene-expression response in adult rat islets

We examined the same eight representative genes in adult primary rat islets to illustrate pathway-specific responses across the 27-gene panel **(Fig. 4F)**. These included *Egln3* for HIF-associated oxygen sensing, *Ddit4* for stress signaling, *Slc2a1, Ldha, Pdk1, Aldoa,* and *Eno1* for metabolic adaptation, and *Vegfa* for vascular adaptation. Across these genes, adult rat islets showed clear responses not only at 1% O_2_ but also at 5% O_2_. *Egln3* increased most strongly at 1% O_2_, while several metabolic and vascular-response genes, including *Slc2a1, Ldha, Pdk1, Vegfa, Aldoa,* and *Eno1*, showed higher expression at both 1% and 5% O_2_ relative to 18% O_2_ at selected timepoints. Overall, these representative profiles supported the heatmap and PCA findings and showed that 5% O_2_ frequently remained distinct from 18% O_2_ in adult rat islets. Complete comparisons are provided in **Supplementary Table 11**, and the full 27-gene profiles are shown in **Supplementary Fig. S4**.

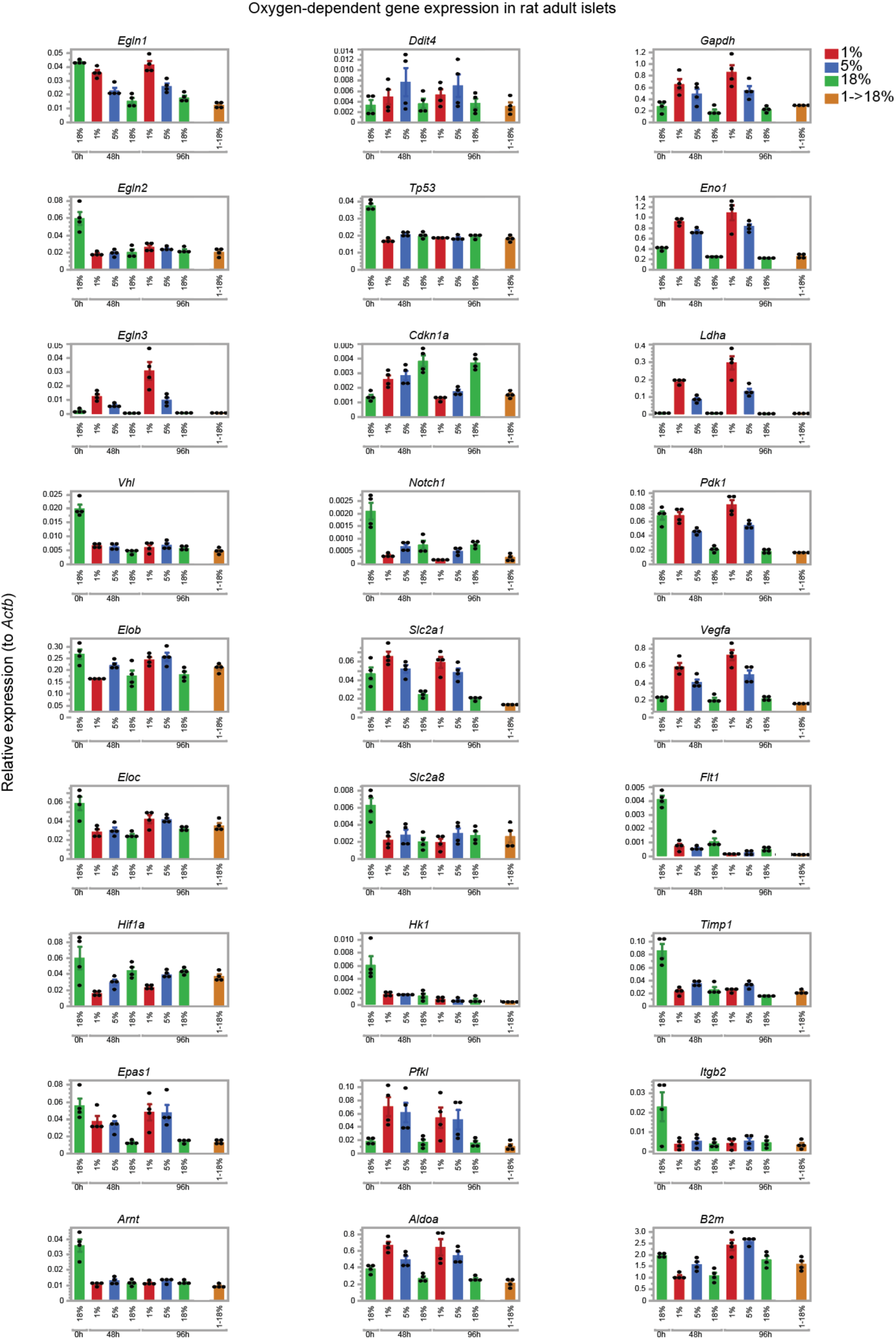

### Distinct oxygen- and time-dependent patterns of islet viability and oxygen sensing

We summarized the oxygen- and time-dependent changes in islet viability, including viability and morphology, alongside oxygen sensing assessed through hypoxia-responsive gene expression **(Fig. 5)**. Adult primary rat islets exhibited progressive loss of viability at 1% O_2_, which occurred predominantly within the first 48 h, whereas overall viability was preserved at 5% and 18% O_2_ through 96 h. Hypoxia-responsive gene-expression patterns persisted at both 1% and 5% O_2_ at 48 and 96 h and remained distinct from those at 18% O_2_. Thus, 5% O_2_ supported short-term islet survival while eliciting a sustained hypoxia-responsive signature, indicating that oxygen-sensing responses occurred without marked loss of overall viability. Following reoxygenation from 1% to 18% O_2_, viability and estimated viable mass did not fully recover to the levels observed under continuous culture at 18% O_2_, whereas gene-expression profiles shifted toward a normoxia-like signature. Together, these findings highlight distinct phenotypic and gene-expression responses to both sustained hypoxia and reoxygenation.

**Figure 5.**
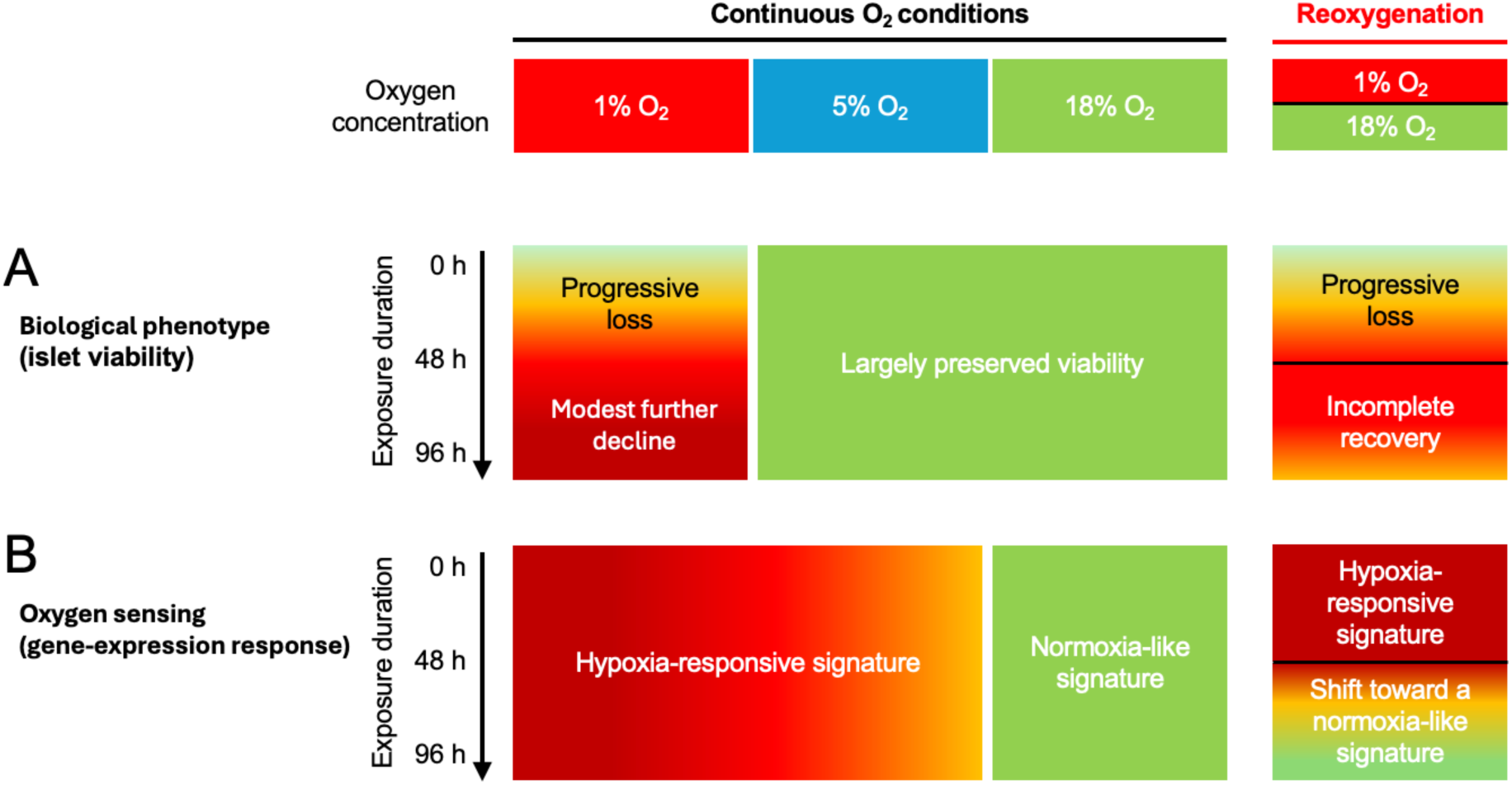
Schematic summary of oxygen- and time-dependent patterns of islet viability and oxygen sensing in adult primary rat islets. Columns indicate continuous culture at 1%, 5%, or 18% O_2_ and reoxygenation from 1% to 18% O_2_ after 48 h. Exposure duration increases downward from 0 to 96 h. **(A)** Overall islet viability. Green indicates largely preserved viability, whereas yellow-to-red shading indicates increasing viability loss. **(B)** Oxygen sensing assessed through gene-expression responses. Red-to-orange shading indicates hypoxia-responsive signatures, whereas green indicates a normoxia-like signature. In both panels, horizontal black lines in the reoxygenation column mark the transition at 48 h. Colors and gradients represent conceptual response patterns rather than quantitative scales or continuously measured changes.

## DISCUSSION

Adult primary rat islets showed substantial vulnerability to severe hypoxia, with viability loss occurring predominantly within the first 48 h at 1% O_2_, whereas overall viability remained largely preserved at 5% and 18% O_2_. Notably, these phenotypic responses differed from the oxygen-sensing responses assessed through gene expression: both 1% and 5% O_2_ elicited sustained hypoxia-responsive signatures that were distinct from those at 18% O_2_. Reoxygenation after an initial 48 h at 1% O_2_ did not fully restore viability to the levels observed under constant 18% O_2_, although it shifted gene-expression profiles toward a normoxia-like signature. Together, these findings demonstrate a separation between oxygen-sensing responses and phenotypic injury. Extending our previous 48-h pO_2__survival analysis (13), the present experiments demonstrated acute islet injury during severe hypoxia and incomplete recovery following reoxygenation. These findings are consistent with early post-transplant islet vulnerability and persistent injury following hypoxia/reoxygenation (19, 20), which highlights the importance of minimizing the severity and duration of early graft hypoxia. Temporary oxygen supplementation may bridge the period before effective revascularization, consistent with improved engraftment following short-term post-transplant oxygenation (21, 22). Interestingly, culture at 5% O_2_ supported overall viability, mass, and estimated viable mass at levels comparable to those at 18% O_2_, suggesting a potential target for short-term post-transplant graft oxygenation. With the oxygen-permeable plate bottom, the 5% O_2_ condition provided a pO_2_ of approximately 40 mmHg, close to oxygen levels reported in several tissues (23). Including this physiologically relevant oxygen condition is a strength of our study, enabling in vitro assessment under conditions that approximate the in vivo tissue environment and informing clinical translation. However, several factors should be considered when applying this finding to transplantation.

Larger islets remained vulnerable at 5% O_2_, and preserved overall viability does not establish sufficient oxygenation for endocrine function or longer-term survival.

Furthermore, achieving this oxygen level at each islet surface requires attention to graft architecture; islet aggregation and dense packing limit oxygen diffusion, causing intragraft oxygen depletion (3, 24, 25). Extrahepatic graft design should optimize islet size, packing density, transplantation volume, and oxygen delivery to maintain adequate oxygenation at individual islet surfaces (4, 26, 27).

Our current study used an oxygen-permeable culture platform to establish well-controlled oxygen conditions, followed by extensive size-stratified analysis of more than 6,000 individual islets. This approach enabled detailed assessment of size-dependent islet responses to oxygen availability. For example, although overall viability did not differ significantly between 5% and 18% O_2_, size-stratified analysis revealed significantly lower viability at 5% O_2_ among large islets. These findings highlight size-dependent vulnerability that was not apparent in the overall analysis and are consistent with our previous study showing that a high proportion of large islets in transplant preparations adversely affected transplantation outcomes (2, 15, 28, 29).

Our study linked phenotypic responses to hypoxia-associated gene-expression signatures across different O_2_ conditions. Although overall viability remained largely preserved at 5% O_2_, detailed gene-expression analysis using PCA revealed that the profile at 5% O_2_ more closely resembled that at 1% than at 18% O_2_. These findings suggest that islets sensed 5% O_2_ as a hypoxic environment despite maintaining overall viability. Previous studies demonstrated rapid oxygen-dependent impairment of islet function (1, 30). Moreover, hypoxia-associated gene signatures in isolated islets can predict impaired graft function (16, 31). Thus, the sustained hypoxia-responsive signature at 5% O_2_ increases the risk of functional compromise despite preserved survival. This distinction necessitates post-transplant strategies that prioritize maintaining islet survival through temporary oxygen support until effective revascularization (4). The 5% O_2_ condition largely preserved short-term overall viability and may therefore help define the minimum oxygen level needed during this period, with the potential for later functional recovery as oxygenation improves. The shift toward a normoxia-like gene-expression signature following reoxygenation is consistent with the possibility of functional recovery, although this requires direct evaluation.

Several limitations should be considered. First, this study prioritized examination of islet survival, and endocrine function was not directly evaluated under each O_2_ condition.

Oxygen-dependent impairment of islet function has been well documented (1, 30). This distinction between survival and function is also relevant to stem cell-derived beta cells, as prolonged reduced-O_2_ exposure can impair beta-cell identity and function even when survival remains relatively preserved (32). Second, this study used adult primary rat islets. Our previous studies demonstrated similar hypoxia susceptibility in adult rat and human islets (13, 33). We therefore anticipate broadly similar responses in adult islets across the species, although direct confirmation is required. We also demonstrated substantial variability in hypoxia resistance among deceased human donors and among individual islets within the same preparation. Third, several technical limitations should be considered. Mass recovery was estimated from two-dimensional images rather than directly quantified using measures such as DNA content (34). Image-based area measurements have been shown to be useful for estimating islet volume (35, 36), but may not fully capture changes in three-dimensional tissue mass or cell number. On the other hand, this approach provides a simple, nondestructive assessment that preserves islets for subsequent assays (35, 37). It also permits repeated measurements of the same preparation at different timepoints, allowing normalization to its own pre-culture value. This longitudinal assessment of the same preparation is a strength of the method despite the limitations of area-based mass estimation. Another technical limitation is the use of a targeted gene panel designed to relate the observed phenotypes to established hypoxia-responsive pathways, which did not define the genome-wide molecular response to reduced O_2_. Previous studies describing broader transcriptomic, proteomic, and metabolic changes associated with stem cell-derived islet maturation and hypoxia responses provide useful context (38).

Overall, our in vitro findings characterized the O_2_-dependent phenotypic and gene-expression responses of adult primary rat islets, providing translational insight into oxygen-support strategies during the early post-transplant hypoxic period.

## SUPPLEMENTAL MATERIAL

Supplemental Figs. S1–S4 and Tables S1–S11

## DATA AVAILABILITY

Data are available upon request.

## ACKNOWLEDGMENTS

N/A

## AUTHOR CONTRIBUTIONS

H. Komatsu conceived and designed the research; H. Kato, H.N.Z., N.E., H.M., and H. Komatsu performed experiments; H. Kato, H.N.Z., N.E., and H. Komatsu analyzed data; H. Kato, H.N.Z., N.E., H.M., H.T.K., and H. Komatsu interpreted the experimental results; H.N.Z. and H. Komatsu prepared figures and drafted the manuscript; H.N.Z., H.T.K., and H. Komatsu edited and revised the manuscript; all authors approved the final version of the manuscript.

## SUPPLEMENTARY FIGURE LEGENDS

**Supplementary Figure S1. Overall size distribution of adult rat islets.**

**(A)** Overall distribution of islet diameters across all analyzed oxygen-exposure conditions and time points.

**(B)** Islet-size distributions shown separately for batches #1–4. Histograms show the number of islets within each diameter range. Islet size is expressed as diameter (µm).

**Supplementary Figure S2. Distribution of viability measurements in adult rat islets across four preparations.**

**(A)** Violin plots showing the distribution of individual islet-level viability measurements across all islet sizes, combining data from four independent adult rat islet preparations. A total of 6,699 islets were analyzed. Conditions include pre-hypoxia (0 h); continuous culture at 1%, 5%, or 18% O_2_ for 48 or 96 h; and reoxygenation consisting of 48 h at 1% O_2_ followed by 48 h at 18% O_2_. Each violin represents the distribution of viability values for all islets analyzed within the indicated condition.

**(B)** The same islet-level viability data shown separately for each of the four independent preparations (batches #1–4). Viability is expressed as the percentage of live cells within each islet. The numbers of islets analyzed under each condition and within each preparation are provided in **Supplementary Table 2**.

**Supplementary Figure S3. PC1 loading coefficients for oxygen-responsive gene-expression profiles in adult primary rat islets.**

PC1 loading coefficients derived from PCA of the common 27-gene panel in adult primary rat islets after **(A)** 48 h and **(B)** 96 h of continuous culture at 1%, 5%, or 18% O_2_. Genes are ordered according to the magnitude and direction of their contribution to PC1 within each analysis. Positive and negative loading directions indicate opposite contributions to the PC1 axis and do not by themselves represent gene upregulation or downregulation.

**Supplementary Figure S4. Oxygen- and time-dependent expression profiles of the full 27-gene panel in adult primary rat islets.**

Relative gene expression before hypoxic exposure (0 h), after 48 or 96 h of culture at 1%, 5%, or 18% O_2_, and following reoxygenation from 1% to 18% O_2_ after 48 h. Expression values were normalized to *Actb*. Each plot represents one gene, with an independently scaled y-axis. Colors indicate oxygen conditions: red, 1% O_2_; blue, 5% O_2_; green, 18% O_2_; and orange, reoxygenation. Black dots represent individual samples from four independent preparations. Bars represent means, and error bars indicate SEM. Black dots represent individual samples from four independent preparations.

